# Genomic Adaptation of an Autochthonous Cider Yeast Strain to Buckwheat and Barley Wort Under Stressful Brewing Conditions

**DOI:** 10.1101/2025.05.14.654042

**Authors:** Martina Podgoršek, Katja Doberšek, Maja Paš, Miha Tome, Miha Ocvirk, Uroš Petrovič, Iztok Jože Košir, Neža Čadež

## Abstract

Growing consumer demand for specialty beers with unique flavors and enhanced nutritional properties is driving the development of novel, high-performance industrial yeasts. However, the genetic diversity of beer yeast strains is limited. Traditional spontaneous fermentations are a rich source of new strains that are well adapted to fermentative environments but lack the ability to efficiently convert maltose-based substrates that are rich in polyphenols (e.g., buckwheat wort) or maltotriose-rich substrates (e.g., barley wort). To simulate the selection pressure exerted on beer yeasts during domestication, we used adaptive laboratory evolution (ALE) to yield cider yeast *Saccharomyces cerevisiae* that can efficiently convert buckwheat and barley wort into beer. To this end, 30 serial transfers of yeast biomass were conducted in high-pressure fermenters simulating industrial-scale stress conditions. This approach resulted in efficient maltose conversion in buckwheat wort and improved maltotriose conversion in barley wort. Three evolved clones from each evolutionary experiment were sequenced using short-read technology and aligned to the chromosome-level assembly of the ancestral cider strain. We observed pronounced genomic changes, including near-complete loss-of-heterozygosity, novel single-nucleotide mutations, and chromosomal aberrations resulting in altered chromosome copy numbers or segmental duplications. Additionally, the clones adapted to buckwheat wort were respiratory-deficient, either lacking or having impaired mitochondrial DNA, whereas clones adapted to barley wort retained a truncated mitochondrial genome. These genetic changes mirror hallmarks of beer yeast domestication and were also reflected phenotypically, including loss of sporulation capacity, decreased fitness under non-brewing conditions, and altered production of aromatic compounds.

**IMPORTANCE:** Consumer demand for specialty beers with distinctive flavors and nutritional value is growing and highlights the need for novel, high-performance beer yeasts adapted to stressful industrial conditions. This study demonstrates how adaptive laboratory evolution can be used to domesticate non-traditional yeasts, enabling efficient fermentation of alternative substrates, such as buckwheat and barley worts. The evolved strains not only improved sugar utilization under industrial conditions but also acquired genomic and phenotypic traits characteristic of domesticated beer yeasts. These findings demonstrate a viable strategy for expanding the functional diversity of brewing yeasts and support innovation in craft beer production.

## INTRODUCTION

The popularity of beer in recent years has led to the continued growth of the craft beer industry, whose competitiveness is driven by a growing interest in new and unique flavors. One of the possible alternatives for new beer types are alternative raw materials such as buckwheat (*Fagopyrum esculentum*), a pseudocereal and melliferous crop (1, 2). Buckwheat is regaining interest as functional food due to its nutritional properties. It is naturally gluten-free, which makes it an excellent option for brewing beer for individuals with celiac disease or gluten intolerance. It is also rich in minerals and bioactive compounds such as runin and quercetin (3, 4), which makes it interesting as a phytonutrient-rich food with potential health benefits.

The introduction of buckwheat into brewing may present some technical challenges, such as haziness in the finished beer due to the high content of polyphenols (1), the need to optimize mashing to exploit the malting ability of grains (5), or alternatively, the use of exogenous enzymes that result in similar values of free-amino nitrogen (FAN) and pH compared to barley wort (2, 6, 7). The final product may have a distinct flavor profile that is more similar to wheat than barley beer (1, 8).

Brewing alternatives are largely driven by consumer demand for a wider range of beer styles, and this has coincided with our increasing understanding of the importance of yeasts in determining the character of beer (9). Recent findings on the population structure of beer yeasts have shown that they are highly influenced by the brewing technology used and their geographic origin (10, 11), highlighting the importance of genomic adaptation for industrially relevant traits. The importance of selecting a yeast strain that is adapted to a new substrate when introducing a new brewing style, such as buckwheat beer, has been substantiated by Deželak et al. (2, 12). They found that the benchmark lager yeast TUM 34/70 had poor performance in buckwheat wort compared to barley wort fermentations.

Therefore, we report here the selection of an autochthonous *Saccharomyces cerevisiae* strain from the complex microbiota of spontaneously fermented traditional foods such as Slovenian cider and its adaptation to a specific beer type by adaptive laboratory evolution (ALE). After 30 successive fermentations under stressful conditions mimicking industrial-scale fermentation, we characterized the genetic basis of adaptations of clones to buckwheat wort compared to barley wort. Furthermore, we evaluated the fitness of these clones in laboratory fermentations and compared their profiles of flavor compounds with those of their ancestor strain. These results provide insight into the genetic mechanisms underlying yeast adaptation to different brewing conditions and may have implications for the development of improved strains for industrial-scale fermentation.

## RESULTS AND DISCUSSION

### Population structure of Saccharomyces yeasts isolated from cider

Considering the limited phenotypic plasticity of lager yeasts for the production of beers from alternative substrates (10), we isolated strains of *Saccharomyces* spp. yeasts from final stages of artisanal cider fermentations at four distinct farms (farm A-D) in Slovenia that never introduced a starter culture. Several *Saccharomyces* colony types were isolated from countable dilution plates and de-replicated using repetitive palindromic element (rep)-PCR fingerprinting (data not shown). From these strains (listed in <u>Supp. Table S1)</u>, 12 genetically distinct *Saccharomyces* spp. strains were subjected to short-read sequencing and identified by mapping the reads to *Saccharomyces* reference genomes (13). As expected, different *Saccharomyces* species and strains coexisted within a single fermentation vessel (Fig. 1). A cryotolerant species of *S. uvarum* with an introgressed genome of *S. eubayanus* was predominant, followed by a hybrid between *S. cerevisiae* and *S. kudriavzevii* (*S.cer* × *S.kud* hybrid) and “pure” *S. cerevisiae*. At one of the farms (farm C), *S. paradoxus* with short introgressions from *S. cerevisiae* was the predominant *Saccharomyces* species, which is rare in domesticated environments (14).

**FIG 1.**
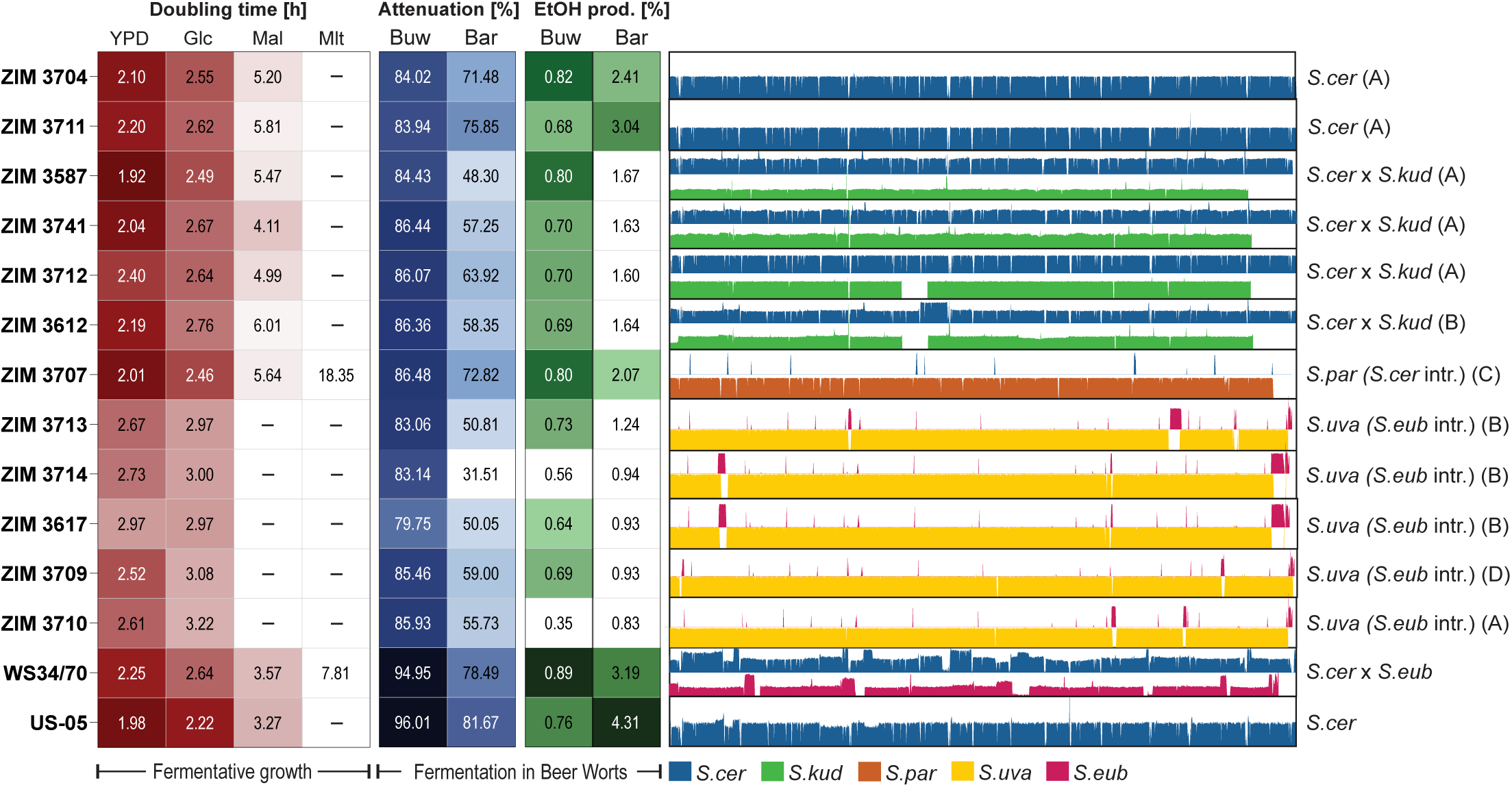
Fermentative performance traits and genomic structure of autochthonous *Saccharomyces* cider strains. Trait heatmaps of *Saccharomyces* cider strain doubling time [h] during fermentative growth in YPD and YNB with glucose (Glc), maltose (Mal), and maltotriose (Mlt) as the sole carbon source and sugar consumption and ethanol production [both in %] in buckwheat (Buw) and barley (Bar) worts. *S. cerevisiae* (*S.cer*), *S. kudriavzevii* (*S.kud*), *S. paradoxus* (*S.par*), *S. uvarum* (*S.uva*), and *S. eubayanus* (*S.eub*) genomic contributions and introgressions (intr.) are shown. The farms (identified by letters A-D) from which strains originated are specified in brackets. The lager yeast strain Weihenstephan WS34/70 and ale yeast SafAle^TM^ US-05 were used as controls.

To examine the evolutionary history of the predominating *Saccharomyces* strains from cider, phylogenetic trees based on SNPs of protein-coding genes were constructed (<u>Fig. S1,</u> <u>A and B</u>). The *S. cerevisiae* genomes and *S.cer* subgenome contributing to the *S.cer* × *S.kud* hybrids formed a monophyletic group within the Wine lineage, known for its genetic features indicating adaptation to stresses in alcoholic fermentation processes (15). Furthermore, the basal position of subgenomes in *S.cer* × *S.kud* hybrids (ZIM 3712, ZIM 3741, ZIM 3578) relative to pure *S. cerevisiae* cider strains (ZIM 3711, ZIM 3704) suggests the independent emergence of interspecies hybrids within this specific cider-producing niche. Similarly, *S. uvarum* strains (<u>Fig. S1B</u>) showing multiple introgressions from *S. eubayanus* were placed among domesticated strains from cider, wine, and beer of the Holarctic lineage, being closest to a cider strain from Germany (16).

### Phenotypic characteristics of cider strains for buckwheat and barley beer fermentations

A comparative phenotypic evaluation of beer fermentation traits of the 12 strains from cider was conducted in comparison to lager yeast (WS34/70) and ale yeast (US-05) strains. We measured cell doubling time under respiration-limited conditions in media containing simple wort sugars (glucose, maltose, and maltotriose) as sole carbon sources and in rich medium (YPD) and fermentation performance (sugar consumption and ethanol production) in buckwheat and barley worts (Fig. 1). Overall, *S. cerevisiae* or hybrid strains showed slower growth on maltose or produced significantly less alcohol compared to beer yeasts. Only *S. paradoxus* was able to ferment maltotriose as the sole carbon source, albeit at a slow rate. In addition, all *S. uvarum* strains showed poor beer fermentation performance with either pure sugars or complex worts under brewing conditions. As expected, based on the evolutionary adaptation of *S. cerevisiae* to human fermentation processes, *S. cerevisae* ZIM 3704 was selected as an alternative yeast strain for buckwheat beer production. Nevertheless, its ability to ferment maltose in buckwheat wort was moderate, and therefore we conducted the evolutionary adaptation experiment with this strain.

### Genomic characterization of the ancestor cider strain

To determine the genetic basis of the adaptation of the natural *S. cerevisiae* cider strain ZIM 3704 to buckwheat and barley worts, its genome was sequenced using long– and short-read sequencing technologies to obtain a high-quality whole-genome sequence for reference in subsequent analyses (<u>Table S2</u>). We assembled its genome into 16 nuclear chromosomes, a mitochondrial chromosome, and a 2µ plasmid using the wine strain DBVPG 6765 for chromosome-level scaffolding and centromere profiling by the LRSDAY pipeline to correctly assign chromosomal identity (17). Furthermore, we observed a normal distribution of base frequencies at variable sites (<u>Fig. S2)</u> and found uniform sequence coverage (200 ± 38) across all chromosomes (<u>Fig. S3</u>, <u>Table S3</u>), suggesting that the strain ZIM 3704 is euploid diploid. The level of heterozygosity was 0.8 SNPs/kbp across the unevenly distributed heterozygous regions of the chromosomes, which were subdivided by regions of loss-of-heterozygosity (LOH) (Fig. 2A<u>, Table S4</u>). These regions spanned 37% of the genome, and some of the chromosomes had large portions of LOH events (e.g., chr. II, IX, XI, XII, and XIV). Extrachromosomal elements included the mitochondrial DNA and 2µ plasmid. Nonetheless, these genomic characteristics are common to the European/Wine clade of *S. cerevisiae* populations (18), as we confirmed by phylogenetic placement of the cider strain (<u>Fig. S1</u>).

**FIG 2.**
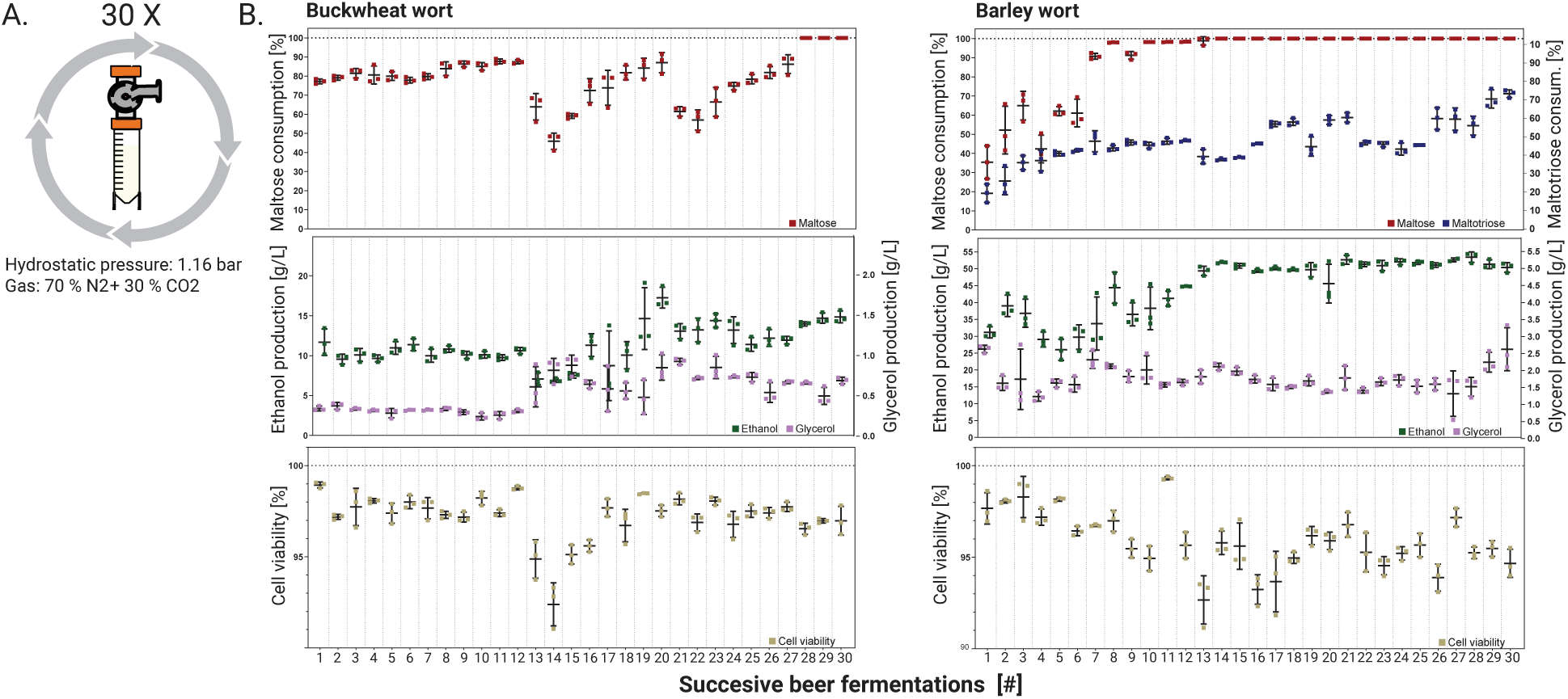
Genome-wide loss-of-heterozygosity and novel mutations in the evolved clones compared to the ancestral cider strain. Chromosome map of the ancestral cider strain, the three clones that evolved in buckwheat wort (A1, A6, and A9), and the three clones that evolved in barley wort (J1, J7, and J9). The following is indicated: heterozygous positions (red bands), new mutations (red triangles), amino acid changes in coding regions (black triangles), and centromeres (grey ovals).

### Adaptation of the cider strain to efficiently ferment sugars of buckwheat and barley worts

Next, we studied the adaptation process, i.e., improved utilization of maltose and maltotriose, of the best performing *S. cerevisiae* ZIM 3704 cider strain to buckwheat and barley worts. For this, we serially transferred yeast biomass in 30 consecutive beer fermentations in an array of high-pressure fermenters simulating the hydrostatic and CO_2_ pressure stress conditions that yeasts are exposed to on an industrial scale (<u>Fig. S4</u>).

During single-batch fermentation, yeast populations grew four and five generations on buckwheat and barley wort, reaching a total of 120 and 150 generations (<u>Table S5</u>), respectively. At each of the 30 serial transfers of yeast biomass, we determined maltose and maltotriose conversion rates, ethanol and glycerol production, and biomass cell viability using methylene blue staining in triplicate (Fig. 3).

**FIG 3.**
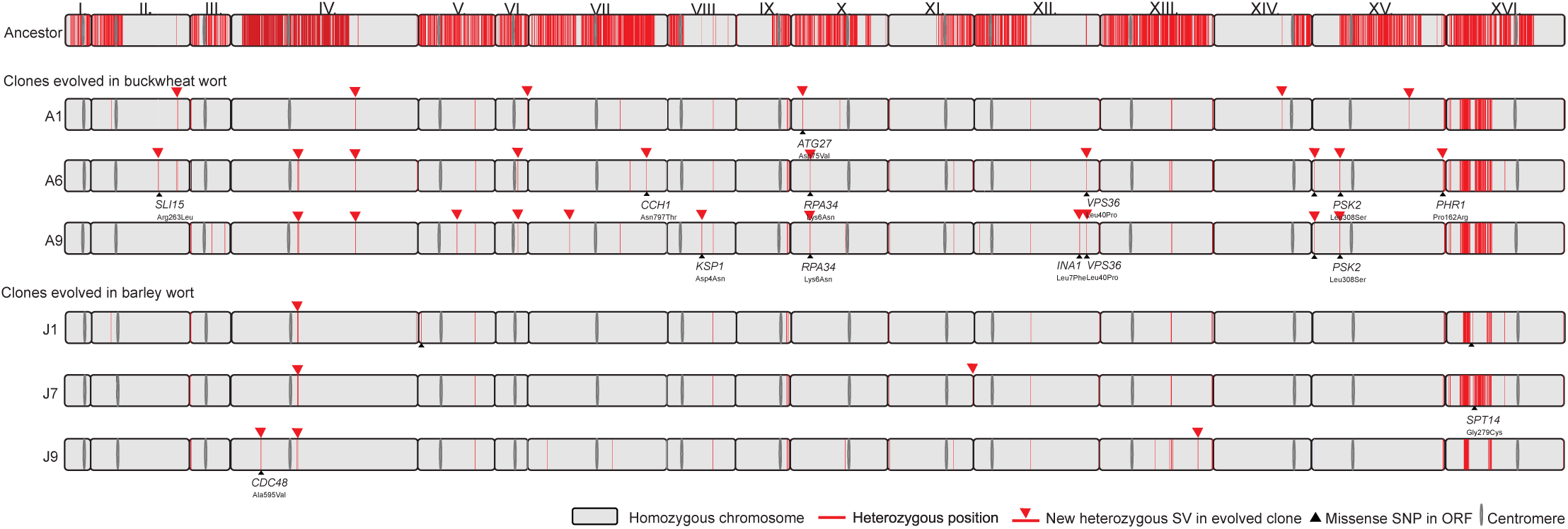
Changes in sugar consumption, product formation, and cell viability during consecutive fermentations of buckwheat and barley worts under stressful conditions. (A) Experimental design using high-pressure fermenters. (B) Scatter plots of maltose and maltotriose consumption (final vs. initial concentration), ethanol and glycerol production, and cell viability (determined by methylene blue staining) for each successive fermentation of buckwheat and barley wort. The horizontal lines represent the mean values of three fermentations.

Through a batch mode of adaptation to utilize maltose in buckwheat wort, maltose consumption generally fluctuated in a reproducible manner in all three replicate batch fermentations. In the first 12 batches, the maltose consumption, ethanol and glycerol production, and cell viability were constant. However, the yeast could convert only approximately 80% of maltose in buckwheat wort. In the 13^th^ fermentation batch, a significant decrease in viability resulted in decreased maltose conversion (from 87 ± 0.8% to 64 ± 5.6%) and increased glycerol production (from 0.31 ± 0.02 g/L to 0.61 ± 0.2 g/L). The latter can be attributed to increased stress conditions due to anaerobiosis, which can lead to increased glycerol production due to reoxidation of NADH formed by assimilatory reactions in mitochondria (19, 20). Most likely, the stressful conditions led to selective advantages for spontaneous mutations that improved stress tolerance in some of the evolved variants. With each batch, the ability to assimilate maltose improved until the 21^st^ batch, in which this ability decreased (from 87 ± 4.3% to 61 ± 2%). However, at this stage, the viability of the evolving variants was in line with previous measurements. From the 27^th^ and until the last, 30^th^ fermentation batch, all maltose in buckwheat wort could be fermented, suggesting that newly evolved variants with improved affinity for maltose (21) displaced existing variants. This “take over” of variants occurred after approximately 108 generations.

The evolutionary adaptation of the cider strain to barley/malt wort was faster than to buckwheat wort, as 100% maltose consumption from the wort had already occurred after the 8^th^ batch (approximately 40 generations). However, the adaptation of the strain to the efficient use of maltotriose was slower and only gradually increased with many fluctuations. In the 30^th^ fermentation batch (approximately 150 generations), the decrease in maltotriose was the highest, reaching 73 ± 1.7% and corresponding to a 2.5-fold improvement. Similar to maltose uptake, the increased maltotriose consumption may be attributed to improved transport capacity (22).

### Genetic changes of evolved clones adapted to buckwheat and barley worts

In total, 30 fermentation batches yielded 132 and 155 generations in buckwheat and barley worts, respectively. From each evolved population in both types of worts, three morphologically distinct clones (<u>Fig. S5</u>) were sequenced using short-read technology and aligned to the ancestral cider strain.

## LOH

We found that almost all heterozygous sites of the ancestral strain were converted to homozygous positions in evolved clones, indicating whole-chromosome LOH in all six clones regardless of wort composition (Fig. 3B). Specifically, during ALE, the clones acquired from 37% of LOH regions of the ancestor to up to 99.2% of LOH in evolved clones, and thus all chromosomes, except chr. XVI, became nearly homozygous. The mechanisms underlying LOH involve either meiotic or mitotic recombination by crossing-over or gene conversion (23, 24). Despite efficient sporulation of the ancestral strain on sporulation media (53%), we did not observe any ascospores in biomass after each re-pitching (results not shown). On this basis, we hypothesized that non-reciprocal exchanges between homologous chromosomes during mitotic growth led to LOH. By using predictive calculations of Dutta et al. (25), we estimated LOH site rates in evolved clones to range from 5.7 × 10^-2^ to 7.4 × 10^-2^ per SNP per division, which is not consistent with the site rates of approximately 10^-4^ to 10^-5^ per SNP per cell division, as is characteristic of mitotic recombination (25, 26). Mitotic recombination events are relatively rare and limited in their length to an average of 7 kb (reviewed by (27)). Thus, we hypothesize that the observed nearly homozygous clones must have undergone also other mechanisms independent of mitotic recombination, such as chromosomal polyploidization and subsequent loss.

### New mutations

Based on the above, we have demonstrated that most of the single-nucleotide mutations in the evolved clones occurred as LOH events (≈ 97%), which most likely unmasked beneficial recessive alleles and conferred specific fitness advantages to the evolving clones in both fermentation environments (27). All newly arisen mutations were heterozygous and had a mean of 40±8 SNPs/clone (Fig. 3B), of which 4.2±2.6 occurred in protein-coding regions of genes (<u>Table S6</u>). Most SNPs were missense variants in buckwheat-wort-adapted clones A6 and A9, with eight and seven mutations, respectively, whereas barley-wort-adapted clones accumulated only two to three missense mutations. For all adaptive mutations, the SnpEff program predicted only moderate fitness effects (<u>Table S6</u>). Nevertheless, most beneficial mutations for a fermentative environment most likely accumulated through LOH mechanisms. However, only a few additional beneficial variants seemingly increased adaptability to the mineral– and polyphenol-rich buckwheat wort.

### Whole-chromosome aneuploidies

As whole-chromosome LOH could be a result of mitotic nondisjunction leading to aneuploidy, we further analyzed the chromosomal aberrations in evolved clones to detect changes in chromosome copy number or segmental duplications. After 30 bottlenecks, the evolved clones gained several chromosomal trisomies and tetrasomies of smaller chromosomes (I, III, V, VI, and IX), which were more pronounced in buckwheat-wort-adapted clones (5–6 events) than in barley-wort-adapted clones (0–4 events) (Fig. 4<u>, Table</u> <u>S7</u>). Our results are consistent with findings that smaller chromosomes are more prone to aneuploidy (28, 29). However, chromosomal size is not the only determinant of aneuploidy, as ploidy change is also an important mechanism for adaptation to stressful environments (30). In our experiment, the stressors were similar to those encountered by brewing yeasts under industrial conditions, such as high hydrostatic pressure, anaerobiosis, low temperature, high osmotic pressure at the beginning, and ethanol stress at the end of fermentations (31), which caused aneuploidy patterns similar to those found in industrial strains (28, 32). Nevertheless, the results indicate that the stringency of the substrate, in this case buckwheat *vs.* barley wort, was the primary factor influencing polyploidisation as a selective advantage.

**FIG 4.**
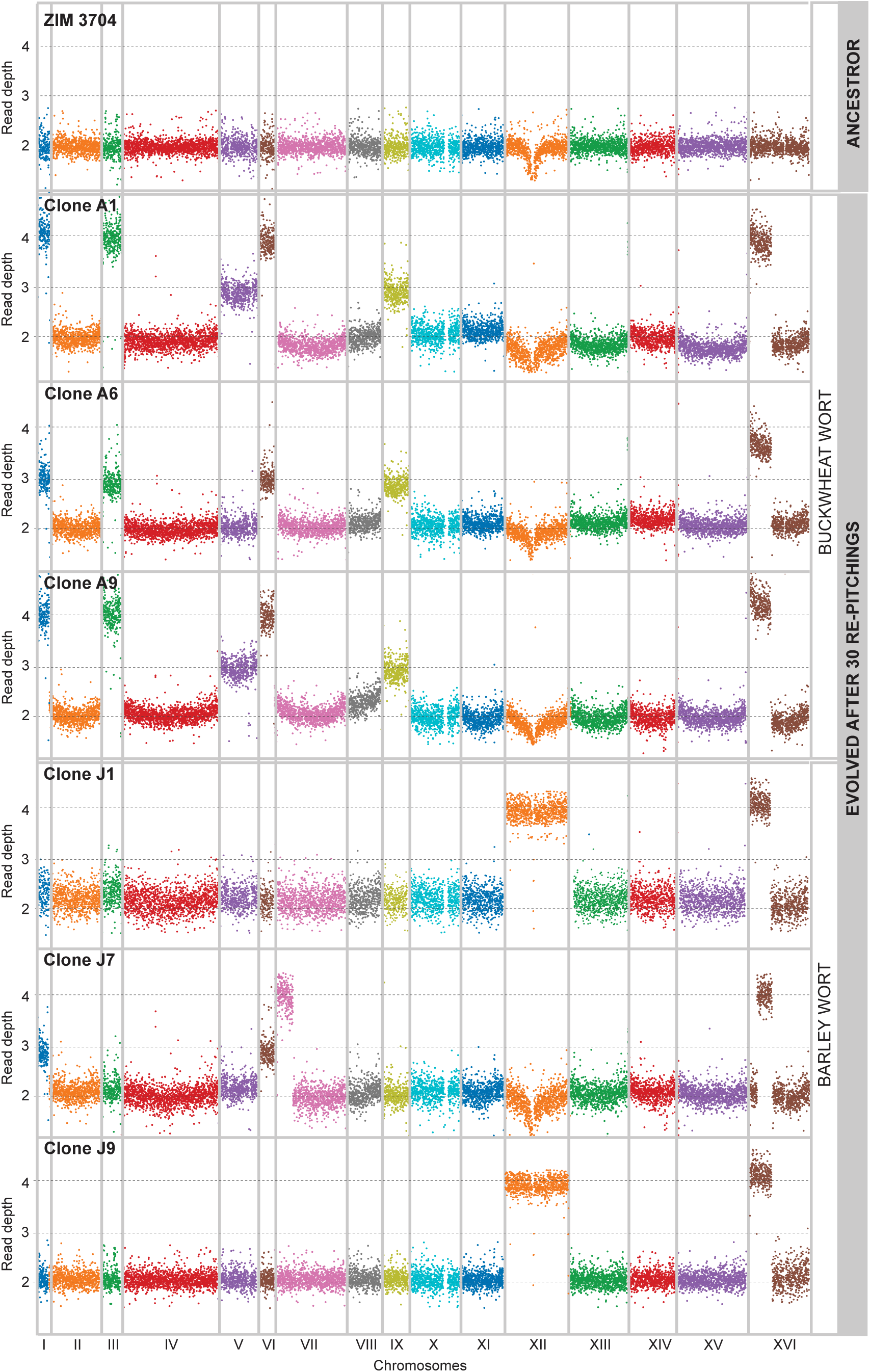
Chromosome copy numbers and large duplications in evolved clones. Genome-wide Manhattan plots predicting ploidy based on read depth of aligned reads for the ancestral cider strain and its evolved clones on buckwheat (A1, A6, and A9) and barley (J1, J7, and J9) worts. Bin size is 100 kb.

### Large segmental aneuploidies

In addition, the large segmental duplications as complex aneuploidies were associated with two larger chromosomes: VII (clone J7) and XVI (all clones). In both cases, in proximity to chromosomal breakpoints, two Ty2 transposable elements arrayed head-to-head at a distance of 220 and 250 kb, respectively (Fig. 5A<u>, Fig. S6</u>). The inverted positions of the Ty elements represent hotspots for the formation of cruciform structures. This can lead to double-strand DNA breaks, which could be further repaired by homology-directed-repair (HDR) (33). This caused a segmental duplication from the left Ty2 element outward almost to the left telomere on one side (≈300 kb) and toward the centromere (≈20 kb) on the other side and a complete deletion of a 14 kb long region for which the ancestor was hemizygous (Fig. 5A). This deletion resulted in a complete loss of 10 genes (<u>Table S8</u>) in all evolved clones.

**FIG 5.**
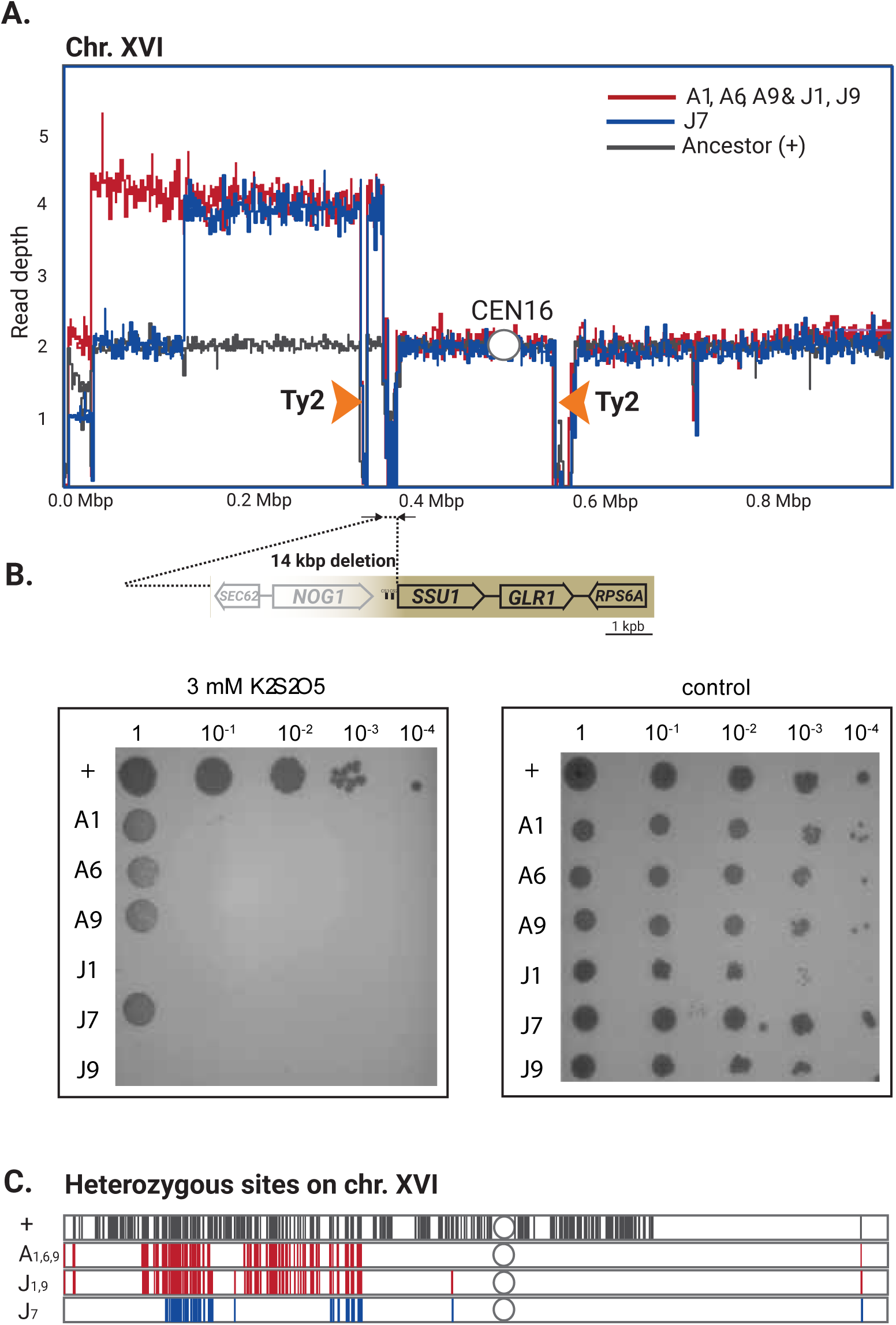
Large segmental aneuploidies from Ty2 elements on chr. XVI result in a deletion of *SSU1* gene promoter causing increased sulfite sensitivity and partial loss of heterozygosity in evolved clones. (A) Read depth profiles showing chr. XVI breakpoints in proximity to head-to-head arrayed Ty2 elements (orange arrows) in evolved clones (red and blue lines) compared to the ancestor strain (black line). (B) A large 14 kb deletion leading to complete loss of 10 genes and a promoter region of the *SSU1* gene causing sensitivity to 3-mM concentrations of K_2_S_2_O_5_ on YPD with 75 mM tartaric acid of evolved clones in comparison to ancestor (+). (C) Partial retainment of the heterozygous region on the left arm of chr. XVI (+, ancestor) in buckwheat (A1, A6, and A9) and barley (J1, J7, and J9) adapted clones, on colored according to the breakpoints (A).

Interestingly, the first gene after the deletion was *SSU1*, which encodes the sulfite efflux pump (Fig. 5B). The activity of Ssu1p increases the excretion of sulfite, which functions as an antioxidant that provides greater flavor stability in beer (34, 35). Even though the *SSU1* gene was retained after these chromosomal rearrangements in evolved clones, the promoter region with the Fzf1 transcriptional factor binding site located 87 bp downstream of the *SSU1* gene was a part of a 14 kbp deletion. Because it has been previously shown that this promoter region has a strong impact on *SSU1* gene expression and, consequently, on the sulfite resistance (36–38) we tested the change in sulfite tolerance of evolved clones in spotting assays using increasing concentrations of K_2_S_2_O_5_ (Fig. 5B<u>, Fig.S7</u>). Consistent with the deletion of the *SSU1* promoter, the evolved clones displayed a significant growth reduction at 3 mM K_2_S_2_O_5,_ whereas the ancestor remained resistant even at higher concentrations (4 mM). Nevertheless, we found that the promoter region of the *SSU1* gene represents a recombination hotspot leading to strain-specific genomic architecture and is one of the factors facilitating the adaptation mechanisms of *S. cerevisiae* to fermentative environments.

Additionally, we found that segmental duplication retained part of the heterozygosity of the parental strain (Fig. 5C) and, based on our findings, we predict the duplication was templated from a homologous region of a chromosome XVI (39). A similar segmental duplication occurred on chr. VII, but only on barley-wort-adapted clone J7 and with a duplicated region induced 160 kb away from a left mirror-oriented Ty2 element pair (<u>Fig.</u> <u>S6</u>). Head-to-head Ty elements in ancestor strain were found only on chr. VII and XVI (<u>Fig.</u> <u>S8</u>)

We found high-rate aneuploidy in most of the buckwheat– and barley-wort-adapted clones. These clones gained the capability to effectively convert maltose in a stressful environment during the evolutionary experiment. However, none of the detected genetic changes are directly related to *MAL* loci. Thus, we predict that LOH together with chromosomal polyploidization might have led to changes in gene dosage and consequently metabolic rewiring.

### Mitochondrial genome

A well-known phenomenon in brewing is the occurrence of respiratory-deficient or “petite” mutants that either lack mitochondrial DNA (ρ^0^phenotype) or are impaired (ρ ^─^ phenotype). These mutants arise spontaneously in response to various stress factors under industrial conditions (40). Since the design of our evolutionary experiment mimics such conditions, we predicted that the small colonies formed by buckwheat-wort-adapted clones (<u>Fig. S5</u>) might be due to their being cytoplasmic “petite”. From *de novo* assemblies of the mitochondrial DNA of the ancestor and evolved clones, we compared their size and gene content after 150 generations under fermentative conditions (Fig. 6A). Of the three buckwheat-wort-adapted clones, clone A1 lost the complete mitochondrial genome (ρ^0^), whereas clones A6 and A9 kept only the *ATP6* gene and reduced their genome from 82 kb to 5 kb. Their impaired respiratory-mediated ATP production was observed, as they formed small (“petite”) colonies (approximately 30% of the ancestor’s colony size) and were unable to grow on non-fermentable carbon source medium. Conversely, the barley-wort-adapted clones J1 and J9 retained most of the canonical protein-coding genes, except the gene encoding apocytochrome b *(COB*), leading to poor growth on non-fermentable carbon sources (Fig. 6B). Furthermore, the mitochondrial genome of clone J7 underwent structural rearrangements, leading to a duplicated and truncated 15S rRNA gene within the *VAR1* and *COX2* genes, which did not affect growth on glycerol (Fig. 6A).

**FIG 6.**
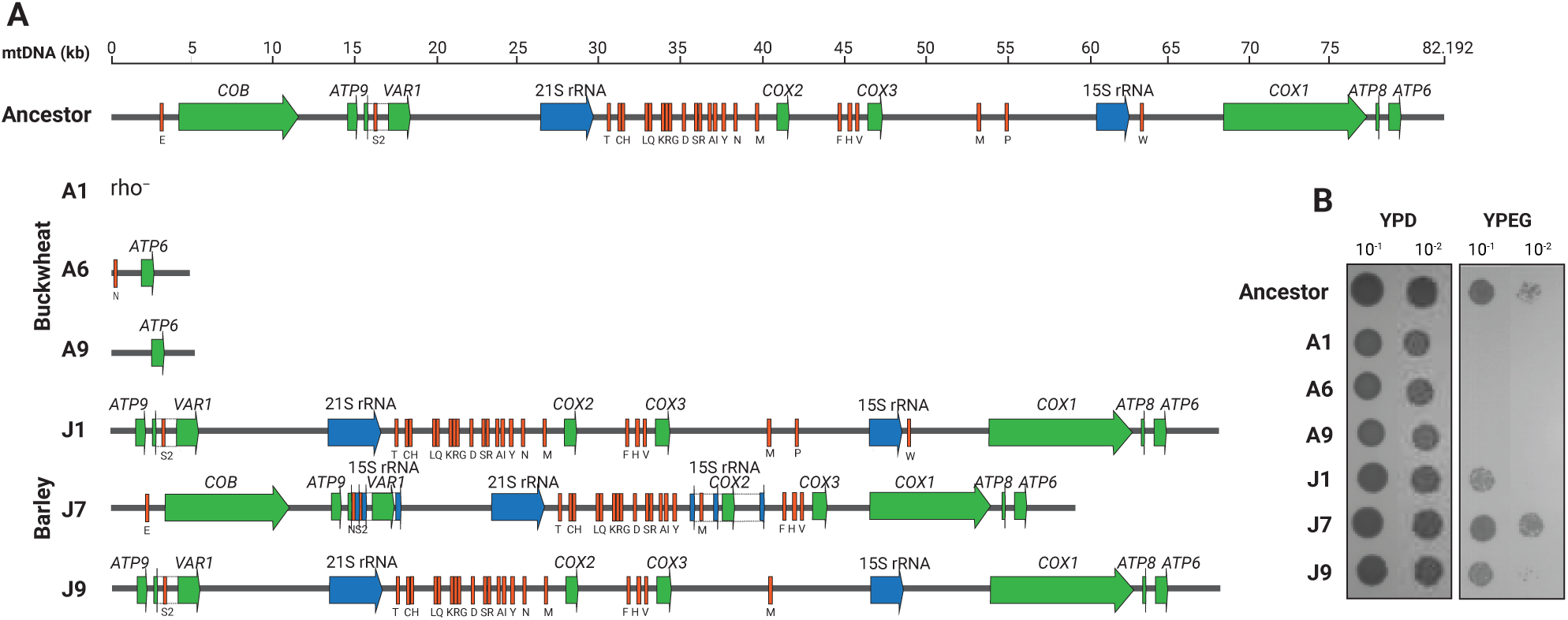
Impact of structural variations in the mitochondrial genomes of the evolved clones on “petite” phenotype. (A) Schematic representation of the mitochondrial genome including protein-coding genes (green), rRNA (blue), and tRNA genes (orange) of clones evolved on buckwheat (A6 and A9) and barley (J1, J7, and J9) worts. Clone A1, rho^─^, lacked mitochondrial DNA. (B) Growth assay on non-fermentable carbon sources glycerol and ethanol (YPEG) in comparison to YPD confirms.

Although it has been generally assumed that the formation of “petite” mutants is caused by many different stress factors during brewery fermentation, strain dependency, and biomass recycling (31, 40, 41), we found that also substrate composition strongly affects “petite” formation (42). During the evolutionary experiment, the nutritionally imbalanced substrate of buckwheat wort caused chromosomal instability that caused polyploidy of some of the chromosomes. This impairment was most likely reflected also in the formation of rho^0^ mutants (43).

### The effects of genetic adaptation on fitness, sexual cycle, and fermentative profile of beer

To understand how the observed genetic changes affect asexual growth, the sexual life cycle, and the ability to produce fermentative aromas, we examined the phenotypes of the adapted clones. Growth rate on a specific substrate is most often used as an indication of yeast fitness (44), and thus we performed growth assays in microtiter plates. Compared to the ancestor strain, clones adapted to barley (J) grew similarly (t-test, p < 0.0001) in buckwheat and barley worts, with a relative fitness of 87% ± 4% and 81% ± 5%, respectively (Fig. 7A). Conversely, the buckwheat-wort-adapted clones (A) grew significantly faster in buckwheat (71% ± 2%) than in barley wort (56% ± 3%), confirming that the genetic changes were selected towards improved fitness for buckwheat wort as a substrate. However, the fitness benefit in clones adapted to specific stress conditions by LOH, aneuploidy, and loss of mitochondrial genes most likely occurred in an additive manner (29, 45). This, however, decreased fitness under different cultivation conditions.

**FIG 7.**
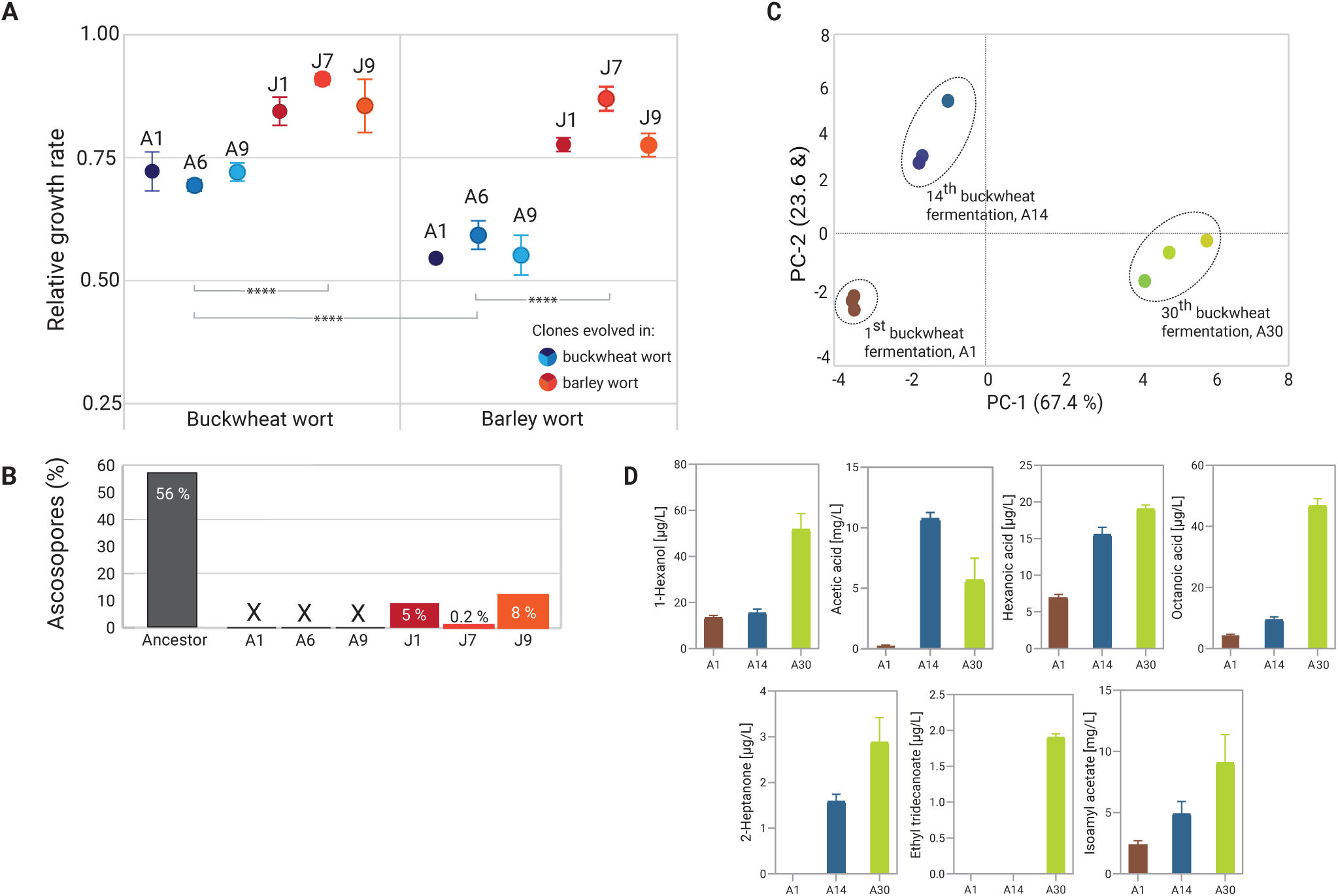
The effects of genomic changes on the phenotypes of the evolved clones and aromatic profiles of buckwheat beers. (A) Relative fitness values of evolved clones compared to the ancestor grown in buckwheat and barley worts in microtiter plates. Error bars show standard deviations from two independent biological replicates. Connecting lines show statistically significant differences between clones (****, p < 0.0001). (B) Sporulation efficiency of evolved clones compared to the ancestor. X, ascospores were not present. (C)PCA score plot of volatile compounds produced by evolving biomass after 1^st^ (A1), 14^th^ (A6) and 30^th^ (A30) successive buckwheat fermentations, each conducted in triplicate. (D) Fermentative volatile compounds that showed statistically significant differences, P<0,005 (one-way ANOVA) in 14^th^ and 30^th^ in comparison to 1^st^ consecutive buckwheat beer fermentation.

Most domesticated strains have an impaired or abolished ability to sporulate (44), and thus we hypothesized that genetic changes acquired during adaptive evolution would result in poor sporulation efficiency, expressed as the percentage of asci relative to vegetative cells (Fig. 7B<u>, Table S9</u>). The ancestral strain had a good sporulation efficiency (55% after 5 days), whereas the buckwheat-wort-adapted clones (A clones) lost sporulation ability due to their respiration deficiency (46). Conversely, the barley-wort-adapted clones (J clones) retained the ability to form tetrads but with a very poor efficiency (0.2–8%). This is in contrast to the findings of Dutta et al. (25), which showed that neither LOH accumulation nor chromosomal instabilities had any effect on fertility, regardless of ploidy level. However, in domesticated *S. cerevisiae* populations, in particular in highly aneuploid beer yeasts, their low sporulation efficiency is partly associated with aneuploidy or with polymorphisms in the promoter region or coding region of *IME1,* a gene encoding a master regulator of meiosis (10, 44). All evolved clones on barley wort had segmental duplications on chromosome XVI, and clone J7 also had a complex aneuploidy on chromosome VII. None of the clones accumulated mutations in the *IME1* gene or its promoter.

Finally, we also explored the effect of evolving populations on fermentative profiles of beer during adaptive evolution. For this, principal-component analysis (PCA) was used to analyze the changes in the aroma profiles of beers after 14^th^, and 30^th^ fermentation batches, which were then compared to the starting beer (Fig. 7C, <u>Table S10</u>). The PCA showed that the distribution of volatile aromas was already clearly separated in the 14^th^ batch by the second principal component (PC2), while PC1 further clearly separated the last, 30^th^ batch. In addition, the fermentative aromas (esters, higher alcohols and medium-chain fatty acids) which differed significantly between the batches (p<0.0001, except isoamyl acetate p = 0.0038, ANOVA) are shown in Fig. 7D. It is noteworthy that in the last, 30^th^ consecutive fermentation, there was a 3 to 20-fold increase in the production of 1-hexanol, acetic acid, hexanoic acid, octanoic acid and isoamyl acetate, with 2-heptanone and ethyl tridecanoate being newly detected in comparison to the 1^st^ batch. Thus, the increasing ability to consume all sugars available in wort of evolved biomass may be the factor for more complex aromatic, fruity, and acidic flavor profile.

## CONCLUSIONS

The growing interest in fermented foods has driven demand for innovative beers that offer consumers new flavors and potential health benefits. The limited genetic diversity of brewing yeasts combined with restrictions on the use of genetically modified organisms in the food and beverage industry has constrained modern approaches to targeted strain improvement. In this study, we used ALE to adapt a strain of *S. cerevisiae*, originally isolated from traditional cider, for the fermentation of both a gluten-free buckwheat wort rich in polyphenols and minerals and conventional barley wort. After 30 consecutive fermentations in a system of high-pressure fermenters designed to simulate stressful industrial conditions, with weekly re-pitching of biomass to fresh wort, we obtained evolved populations that efficiently consumed maltose and produced a more complex aromatic profile of the final product. The evolved clones exhibited genetic changes characteristic of domestication, including loss of heterozygosity, extensive chromosomal or segmental aneuploidy, and rearrangements or loss of mitochondrial DNA. Over the past 200 years, a similar process of continuous “back-slopping” under semi-industrial brewing conditions has led to the emergence of novel yeast phenotypes that are well adapted to the brewing environment. With our experiment, we have recreated the past adaptive evolutionary processes that led to the emergence of new phenotypes.

## MATERIALS AND METHODS

### Yeast strains and maintenance

*Saccharomyces* strains were isolated from the final stages of spontaneously fermented cider collected from remote farms in the low-mountain range in the subalpine region of Koroška in Slovenia. The strains listed in **Supplementary Table 1** are maintained in cryotubes with YPD (Conda) and 20% glycerol at −80 °C in the ZIM Collection of Industrial Microorganisms (www.zim-collection.si). As representative brewing strains, the lager yeast *Saccharomyces pastorianus* TUM 34/70 (Weihenstephan, Freising, Germany) and ale yeast SafAle™ US-05 (Fermentis, Marquette-lez-Lille, France) were used.

### Identification and population genomics of the cider strains

At the genus level, strains were identified by PCR and Sanger sequencing of their ITS region using the primers ITS1 and ITS4 (47). Sequences were assembled in BioNumerics 7.6 (Applied Maths, Sint-Martens-Latem, Belgium) and deposited in GenBank.

The hybrid detection and analysis pipeline spIDer (13) was used to distinguish between pure species, hybrids, and introgressed strains based on genome sequences. For whole-genome sequencing, the TrueSeq DNA PCR Free (350) library was constructed and run on an Illumina NovaSeq instrument at the Macrogen Europe sequencing facility. The resulting short-read data are available in the NCBI Sequence Read Archive (SRA) database under accession number PRJNA818497. Paired-end reads were first trimmed using Trimmomatic v.038 (48) and then aligned using the SpIDer pipeline to concatenated reference genomes of *Saccharomyces* species commonly associated with fermentations, including *S. cerevisiae, S. uvarum, S. eubayanus, S. kudriavzevii*, and *S. paradoxus*.

Next, we inferred the origin of *Saccharomyces* species and their hybrids isolated from cider together with 144 *S. cerevisiae* strains sequenced by Gallone et al. (10) and available in GenBank (BioProject PRJNA323691) and 55 *S. uvarum* strains sequenced by Almeida et al. (16) and Gallone et al. (49). The reads of each strain were mapped either to *S. cerevisiae* S288c (R64-2-1) or *S. uvarum* CBS 7001 (50) reference genomes with Burrows-Wheeler Aligner, BWA v. 0.7.17-r1188, (51) using default parameters. Conversion of SAM to BAM format, sorting, indexing, and quality filtering (MQ<40) were performed using the tools available in the SAMtools package v1.10 (52). Duplicated reads were marked with Picard tools 2.0.1 (https://broadinstitute.github.io/picard/). At each genomic position, a genotype was generated using BCFtools mpileup and subsequently, variants were called using BCFtools call of the SAMtools package. The genetic variants were annotated with SnpEff 5.0e, and only those single nucleotide polymorphisms (SNPs) present in the coding sequences were retained (798,588 and 2,809,113 sites for *S. cerevisiae* and *S. uvarum,* respectively). The variant caller file was converted to alignments for phylogenetic analysis using vcf2phylip y2.8 script (53). The ML phylogenetic tree was constructed in iQtree v.2.2.0.3 using the GTR+ASC model and 1000 bootstrap permutations (54).

### Phenotypic characterization

To assess the fermentation capacity of the strains, experiments were performed in 96-well plates with 200 µL of YNB medium with or without 1% (w/v) glucose, maltose, and maltotriose and supplemented with 3 mg/L antimycin (Sigma-Aldrich, St. Louis, MO, USA) to block respiration (10). The outer wells contained 200 µL of water and were not inoculated. Yeasts for inoculation were pre-cultured in 50 mL conical tubes with 5 mL of YPD medium at 220 rpm and 28 °C for 24 h. Then, cells were centrifuged and resuspended in phosphate-buffered saline. OD_600_ was measured to adjust the final concentration of inoculated cells to OD_600_ ≈ 0.1. Cell growth was monitored every 20 min after 30 s of agitation at 28 °C for 24 h using a microtiter plate reader (Tecan Spark, Männedorf, Switzerland). Mean growth parameters (specific growth rate, lag phase duration, and doubling time) were calculated from two biological replicates, each with three technical replicates, on a single plate using Curveball 0.2.16 (55).

Buckwheat and barley wort fermentations were performed at 14° C in 50 mL (1) conical tubes with a water lock placed on each fermenter. Buckwheat wort (8.3 g/L glucose and 6.6 g/L maltose) was prepared from unmalted buckwheat at the Slovenian Institute of Hop Research and Brewing’s pilot brewery. Enzymes BioproteaseTM NPL, PromaltTM 295TR, Hitempase® 2XL and Bioclucanase® GB (Kerry Food Ingredients Ltd., Naas, Ireland) were added. Barley wort was prepared from unhoped wort concentrate (Döhler, Darmstadt, Germany) diluted to 12°P (10 g/L glucose, 71 g/L maltose, and 25 g/L maltotriose). Extracted iso-alpha acids (30%. Hopfenpflanzerverband Hallertau e.V., Wolnzach, Germany) were added to the wort at a final concentration of 51.5 mg/L. The fermentations were conducted in triplicate, and their progress was monitored daily by measuring the weight loss until no change in weight was observed for 24 h. At the end, the fermentation samples were filtered through 0.45 µm filters, and concentrations of fermentable sugars (glucose, maltose and maltotriose) and ethanol were measured using HPLC. Specifically, a separation module and system interface module liquid chromatograph (Knauer, Berlin, Germany) coupled with a differential refractometer (Knauer, Berlin, Germany) were used. An Aminex HPX-87H organic acid analysis column (300 × 7.8 mm, Bio-Rad) was equilibrated with 5 mM H_2_SO_4_ (Sigma-Aldrich, St. Louis, MA, USA) in water at 40 °C, and samples were eluted with 5 mM H_2_SO_4_ in water at a flow rate of 0.6−1 mL/min.

### Adaptive laboratory evolution with serial transfer in beer fermentations

We evolved the cider yeast strain ZIM 3704 in buckwheat and barley wort as two parallel evolving lines and passed them through 30 bottleneck fermentations, each lasting 7 days (approximately 130-150 generations). Fermentation experiments were conducted in 50 mL conical tubes (TPP, Trasadingen, Switzerland) with silicone tubing connected to the gas pressure control system with an electromagnetic valve that kept the pressure constant at 1.10 ± 0.03 bar to imitate industrial brewery fermentations. The gas mixture supplied to the tubes consisted of 70% N_2_ and 30% CO_2_, achieving the desired CO_2_ partial pressure in the gas phase. The gauge pressure of 1.1 bar in the fermenters mimicked the hydrostatic pressure at the bottom of a 10 m high industrial fermenter at 14° C. The magnetic stirrer was set to 80 rpm to prevent the biomass from settling.

For generation 0 fermentations, yeast cultures were propagated in YPD at 28 °C overnight on a shaker set at 220 rpm. The cell biomass was centrifuged, washed, and resuspended in water to inoculate 35 mL of buckwheat and barley wort at a final concentration of 1×10^7^ cells/mL in triplicate. The fermenters were connected to the gas control system in the incubator set at 14 °C. After 7 days, the fermenters were cooled to 0 °C for 1 h. Biomass was centrifuged at 1500 rpm for 5 min, pooled from three buckwheat and barley fermenters, and used for pitching next generation (1–30) fermentations at a concentration of 1×10^7^ cells/mL. The remaining biomass was analyzed for yeast viability using methylene blue and automatically counted under a microscope using ImageJ software (56) as described by Zupan et al. (57). The wort in the supernatant was analyzed by HPLC for sugar composition and ethanol and glycerol production as described above.

The biomasses after the 30^th^ buckwheat and barley fermentations were diluted to single colonies and streaked onto buckwheat and barley wort agar plates. Three morphologically distinct colonies from each fermentation were isolated and sequenced on Illumina NovaSeq to 200 x depth and deposited in the NCBI SRA under BioProject PRJNA1251027.

### Chromosome-level assembly of the ancestor strain

The *S. cerevisiae* ZIM 3704 strain isolated from cider in Slovenia was sequenced by using a combination of long– and short-read sequencing technologies of PacBio and Illumina, respectively. Sequencing libraries were constructed using the PacBio Sequel Microbial Library Construction (PacBio, California, USA) and TruSeq DNA PCR Free (Illumina, California, USA) kits. The sequencing runs were performed on PacBio Sequel and Illumina NovaSeq instruments at the Macrogen Europe B.V. sequencing facility. The genome is available at DDBJ/ENA/GenBank under the accession JBNCZA000000000.

To generate chromosome-level assembly, PacBio and Illumina reads were processed using the modular computational tool LRSDAY 1.6.0 (17, 58) according to the manual. The assembly file was deposited in the NCBI Genome under BioProject PRJNA1251021. Furthermore, the nuclear and mitochondrial genomes and 2 µ plasmid assembly based on sequence homology comparison to identify protein-coding genes, centromeres, transposable elements of Ty families, and telomere-associated X’– and Y-elements were further automatically annotated using the LRSDAY pipeline. To determine ploidy level of the ancestor strain, we used nQuire v.1.6 software to align short-reads in bam format to an assembled genome, enabling the determination of base frequency distributions at variable sites (59).

### Analysis of loss of hetereozygosity (LOH) and *de novo* mutations

Sequencing reads of the ancestor and evolved clones were mapped to the masked with RepeatMasker(60) chromosome-level of the ancestor assembly genome using BWA-0.7.17 (61), and then converted to bam and sorted with SAMtools v.1.10 (52). Next, duplicated reads were removed by the MarkDuplicates command of Picard tools v. 2.0.1, followed by variant calling using HaplotypeCaller of the Genome Analysis ToolKit (GATK, v.4.1.3.0)(62). The resulting SNPs were filtered by VariantFiltration following best practice of GATK (63): QD (quality by depth) 2.0, FS (Fisher strand bias test score) 60.0, MQ (root mean square of the mapping quality) 40.0, MappingQualityRankSum (mapping quality rank sum test score) 12.5, ReadPosRankSum (read position rank sum test score) −8.0. Indels and SNPs were filtered out, and heterozygous sites were marked by SelectVariants module. The heterozygous sites were manually inspected using Integrative Genomics Viewer 2.12.3 (64). LOH tracts were defined as continuous homozygous regions spanning at least 51 kbp, with each tract containing fewer than 10 interspersed heterozygous positions (65). After masking regions of LOH, the level of heterozygosity was determined as the ratio of heterozygous sites per kbp (18). LOH site rates per clone were calculated as number of sites under LOH divided by the product of genome size and the number of cell divisions (25). The effect of newly arisen mutations in evolved clones was assessed and annotated with SnpEff 5.0e using generated gene annotations of the ancestor strain (66).

### Detection of copy-number and structural variants

The copy-number variations at the chromosomal level were detected based on the depth of coverage of mapped reads to the ancestral genome, using CNVpytor with a bin size of 100 (67). The resulting read depth plots were further validated using custom-made scripts to detect ‘simple’ and ‘complex’ aneuploidies available at https://github.com/SAMtoBAM/aneuploidy_detection, as detailed in O’Donnell et al. (68).

### Analysis of mitochondrial genome architecture and its functionality

We used whole-genome assemblies of evolved clones generated with SPAdes v3.13.1 to identify the mitochondrial scaffold by performing a BLASTn search, with the mitochondrial sequence of the ancestral strain as the query. The MITOS2 wrapper was employed to annotate the mtDNA in FASTA using the refseq63f database to identify candidate sequences for each gene, along with covariance models for the annotation of tRNA and rRNA genes (39). The output in BED file format was then imported into Geneious Prime 2011.1.1 for graphical visualization. The functionality of mitochondria was assessed by spotting 10-fold serial dilutions of overnight cultures pre-grown in liquid YPD medium.

Cultures were suspended in water to a final OD_600_ of 1, then spotted onto YPD and YP plates containing 3% (w/v) glycerol and 3% (w/v) ethanol as the sole carbon sources. The plates were incubated at 28°C for 3 days, and images were acquired using a G:BOX F3 (Syngene)

### Spotting assays

The cultures were pre-grown in liquid YPD medium, washed, and resuspended in water to a final OD₆₀₀ of 1. Starting from an OD₆₀₀ of 1, 10-fold serial dilutions were prepared in sterile water, and 3 µL of each dilution was spotted onto the appropriate plates. Sulfite tolerance was assessed on YPD agar plates containing 75 mM tartaric acid which were overlaid with 0.1 M filter-sterilized potassium metabisulfite solution to achieve the desired final concentrations (1.5 mM, 3 mM and 4 mM) (36). Functionality of mitochondria was assessed by spotting 10-fold serial dilutions onto YPD and YP plates containing 3% (w/v) glycerol and 3% (w/v) ethanol as the sole carbon sources (69). The plates were incubated at 28°C for 3 days, and images were acquired using a G:BOX F3 (Syngene of Synoptics Ltd., Cambridge, UK).

### Fitness of evolved clones

To measure relative fitness of clones after adaptive evolution experiment in comparison to ancestor, the biomass was propagated and prepared as described above and inoculated in final OD_600_ of 0.1 in 200 µL of buckwheat and barley worts with addition of iso-alpha acids in microtiter plates. The growth was monitored after double orbital shaking every 10 minutes in microtiter plate reader (Tecan Spark, Männedorf, Switzerland) for 48 h at 22°C. Two independent replicates were run. The growth kinetics parameters were calculated using PRECOG software (70). Statistical significance was tested using a one-way ANOVA followed by Tukey’s test, implemented in Prism v10.4.0 (GraphPad Inc., La Jolla, CA).

### Sporulation efficiency of evolved clones

To determine the sporulation efficiency of the ancestor and its evolved clones, we followed the protocol described by De Chiara et al. (44), in which, after propagating the cultures in YPD, a 1:50 dilution was inoculated into 10 mL of the pre-sporulation medium YP-acetate (2%) and grown for 48 h at 30°C and 220 rpm. The pre-sporulated cells were transferred to the sporulation medium (2 % potassium acetate) at a ratio of 1:5 in 250 mL Erlenmayer flasks at 23°C and shaken at 220 rpm. To calculate the sporulation efficiency, between 200 and 300 cells were counted under a light microscope at 100x magnification after five days and the ratio of asci to vegetative cells was determined. The buckwheat wort-adapted clones did not grow in a pre-sporulation medium.

### Chemical analysis of beer produced with evolving biomass

Beer samples (10 mL) were transferred into 20 mL GC/MS vials, where 0.5 g NaCl and 0.5 mL of internal standard iso-butanol were added. The samples were analyzed by gas chromatography (Agilent 8890 GC System; Agilent, USA) coupled with mass spectrometry (597BB GC/MS; Agilent, USA) using a system equipped with an autosampler (PAL RSI 120). Agilent MSD ChemStation Enhanced Data Analysis (Rev. F.01.036.2357) was used for data analysis. Volatile compounds in the sample headspace were extracted at 50 °C and 250 rpm for 30 min. A (5% phenyl)-methylpolysiloxane capillary column (30 m length × 250 μm i.d. × 0.25 μm film thickness, Agilent, USA) using He (≥99.999 % purity) as the carrier gas was used to separate the compounds. GC program was set as follows: Initial temperature of 40 °C held for 3 min, increased to 50-100 °C at 6 °C/min held for 2 min, raised to 100-220 ◦C at 10 °C/min, held for 3 min, raised to 220-250 ◦C at 22.5 °C/min, held for 5 min. Mass spectra were recorded in electron ionization (EI) mode at 70 eV with an ion source temperature of 230 °C and a quadrupole temperature of 150 °C. The results were expressed as the ratio between the peak area of the sample and the peak area of the internal standard (IS), multiplied by the IS concentration. Statistical differences were obtained by Prism v10.4.0 (GraphPad Inc., La Jolla, CA) using a one-way ANOVA.

## ACKNOWLEDGMENTS

The authors thank Saška Košir, Eva Urbančič and Lidija Hladnik for technical assistance and Aleš Ogrinec for modelling anaerobic system of fermenters. We thank dr. Eva Lasič for editing and reviewing a draft of this manuscript. We acknowledge Gašper Žun for his valuable comments. Nataša Kočar Mlinarič and Matej Oset formerly of Laško-Union Brewery are acknowledged for their support.

This work was supported by Slovenian Research and Innovation Agency (L4-8222, J4-4552, P4-0116 and MRIC-UL ZIM, IP-0510) and in part by Laško-Union Brewery (Heineken group).

## REFERENCES

1. Phiarais BPN, Mauch A, Schehl BD, Zarnkow M, Gastl M, Herrmann M, Zannini E, Arendt EK. 2010. Processing of a Top Fermented Beer Brewed from 100% Buckwheat Malt with Sensory and Analytical Characterisation. Journal of the Institute of Brewing 116:265–274.

2. Deželak M, Zarnkow M, Becker T, Košir IJ. 2014. Processing of bottom-fermented gluten-free beer-like beverages based on buckwheat and quinoa malt with chemical and sensory characterization. 120:360–370.

3. Giménez-Bastida JA, Zieliński H. 2015. Buckwheat as a Functional Food and Its Effects on Health. J Agric Food Chem 63:7896–913.

4. Skrabanja V, Kreft I, Golob T, Modic M, et al. 2004. Nutrient Content in Buckwheat Milling Fractions. Cereal Chemistry 81:172–176.

5. Wijngaard HH, Arendt EK. 2006. Optimisation of a Mashing Program for 100% Malted Buckwheat. 112:57–65.

6. Ciocan ME, Salamon RV, Ambrus Á, Codină GG, Chetrariu A, Dabija A. 2023. Use of Unmalted and Malted Buckwheat in Brewing. 13:2199.

7. Cela N, Galgano F, Perretti G, Di Cairano M, Tolve R, Condelli N. 2022. Assessment of brewing attitude of unmalted cereals and pseudocereals for gluten free beer production. Food Chemistry 384:132621.

8. Kiss Z, Vecseri-Hegyes B, Kun-Farkas G, Hoschke Á. 2011. Optimization of malting and mashing processes for the production of gluten-free beers %J Acta Alimentaria. 40:67–78.

9. Gibson B, Geertman J-MA, Hittinger CT, Krogerus K, Libkind D, Louis EJ, Magalhães F, Sampaio JP. 2017. New yeasts—new brews: modern approaches to brewing yeast design and development. FEMS Yeast Research 17.

10. Gallone B, Steensels J, Prahl T, Soriaga L, Saels V, Herrera-Malaver B, Merlevede A, Roncoroni M, Voordeckers K, Miraglia L, Teiling C, Steffy B, Taylor M, Schwartz A, Richardson T, White C, Baele G, Maere S, Verstrepen KJ. 2016. Domestication and Divergence of Saccharomyces cerevisiae Beer Yeasts. Cell 166:1397–1410.e16.

11. Goncalves M, Pontes A, Almeida P, Barbosa R, Serra M, Libkind D, Hutzler M, Goncalves P, Sampaio JP. 2016. Distinct Domestication Trajectories in Top-Fermenting Beer Yeasts and Wine Yeasts. Curr Biol 26:2750–2761.

12. Dezelak M, Gebremariam MM, Cadez N, Zupan J, Raspor P, Zarnkow M, Becker T, Kosir IJ. 2014. The influence of serial repitching of Saccharomyces pastorianus on its karyotype and protein profile during the fermentation of gluten-free buckwheat and quinoa wort. Int J Food Microbiol 185:93–102.

13. Langdon QK, Peris D, Baker EP, Opulente DA, Nguyen HV, Bond U, Goncalves P, Sampaio JP, Libkind D, Hittinger CT. 2019. Fermentation innovation through complex hybridization of wild and domesticated yeasts. Nat Ecol Evol 3:1576–1586.

14. Boynton PJ, Wloch-Salamon D, Landermann D, Stukenbrock EH. 2021. Forest Saccharomyces paradoxus are robust to seasonal biotic and abiotic changes. 11:6604–6619.

15. Basile A, De Pascale F, Bianca F, Rossi A, Frizzarin M, De Bernardini N, Bosaro M, Baldisseri A, Antoniali P, Lopreiato R, Treu L, Campanaro S. 2021. Large-scale sequencing and comparative analysis of oenological Saccharomyces cerevisiae strains supported by nanopore refinement of key genomes. Food Microbiology 97:103753.

16. Almeida P, Goncalves C, Teixeira S, Libkind D, Bontrager M, Masneuf-Pomarede I, Albertin W, Durrens P, Sherman DJ, Marullo P, Hittinger CT, Goncalves P, Sampaio JP. 2014. A Gondwanan imprint on global diversity and domestication of wine and cider yeast Saccharomyces uvarum. Nat Commun 5:4044.

17. Yue JX, Liti G. 2018. Long-read sequencing data analysis for yeasts. Nat Protoc 13:1213–1231.

18. Peter J, De Chiara M, Friedrich A, Yue JX, Pflieger D, Bergstrom A, Sigwalt A, Barre B, Freel K, Llored A, Cruaud C, Labadie K, Aury JM, Istace B, Lebrigand K, Barbry P, Engelen S, Lemainque A, Wincker P, Liti G, Schacherer J. 2018. Genome evolution across 1,011 Saccharomyces cerevisiae isolates. Nature 556:339–344.

19. Bakker BM, Overkamp KM, van Maris AJ, Kötter P, Luttik MA, van Dijken JP, Pronk JT. 2001. Stoichiometry and compartmentation of NADH metabolism in Saccharomyces cerevisiae. FEMS Microbiol Rev 25:15–37.

20. Nissen TL, Schulze U, Nielsen J, Villadsen J. 1997. Flux Distributions in Anaerobic, Glucose-Limited Continuous Cultures of Saccharomyces Cerevisiae. 143:203–218.

21. Jansen ML, Daran-Lapujade P, de Winde JH, Piper MD, Pronk JT. 2004. Prolonged maltose-limited cultivation of Saccharomyces cerevisiae selects for cells with improved maltose affinity and hypersensitivity. Appl Environ Microbiol 70:1956–63.

22. Brickwedde A, van den Broek M, Geertman J-MA, Magalhães F, Kuijpers NGA, Gibson B, Pronk JT, Daran J-MG. 2017. Evolutionary Engineering in Chemostat Cultures for Improved Maltotriose Fermentation Kinetics in Saccharomyces pastorianus Lager Brewing Yeast. Frontiers in Microbiology 8.

23. Laureau R, Loeillet S, Salinas F, Bergström A, Legoix-Né P, Liti G, Nicolas A. 2016. Extensive Recombination of a Yeast Diploid Hybrid through Meiotic Reversion. PLOS Genetics 12:e1005781.

24. Dutta A, Lin G, Pankajam AV, Chakraborty P, Bhat N, Steinmetz LM, Nishant KT. 2017. Genome Dynamics of Hybrid Saccharomyces cerevisiae During Vegetative and Meiotic Divisions. G3 Genes|Genomes|Genetics 7:3669–3679.

25. Dutta A, Dutreux F, Schacherer J. 2021. Loss of heterozygosity results in rapid but variable genome homogenization across yeast genetic backgrounds. Elife 10.

26. Sui Y, Qi L, Wu J-K, Wen X-P, Tang X-X, Ma Z-J, Wu X-C, Zhang K, Kokoska RJ, Zheng D-Q, Petes TD. 2020. Genome-wide mapping of spontaneous genetic alterations in diploid yeast cells. 117:28191–28200.

27. Smukowski Heil C. 2023. Loss of Heterozygosity and Its Importance in Evolution. Journal of Molecular Evolution 91:369–377.

28. Gilchrist C, Stelkens R. 2019. Aneuploidy in yeast: Segregation error or adaptation mechanism? Yeast 36:525–539.

29. Sharp NP, Sandell L, James CG, Otto SP. 2018. The genome-wide rate and spectrum of spontaneous mutations differ between haploid and diploid yeast. Proc Natl Acad Sci U S A 115:E5046–E5055.

30. Linder RA, Zabanavar B, Majumder A, Hoang HC, Delgado VG, Tran R, La VT, Leemans SW, Long AD. 2022. Adaptation in Outbred Sexual Yeast is Repeatable, Polygenic and Favors Rare Haplotypes. Mol Biol Evol 39.

31. Gibson BR, Lawrence SJ, Leclaire JP, Powell CD, Smart KA. 2007. Yeast responses to stresses associated with industrial brewery handling. FEMS Microbiol Rev 31:535–69.

32. Scopel EFC, Hose J, Bensasson D, Gasch AP. 2021. Genetic variation in aneuploidy prevalence and tolerance across Saccharomyces cerevisiae lineages. Genetics 217.

33. Lemoine FJ, Degtyareva NP, Lobachev K, Petes TD. 2005. Chromosomal translocations in yeast induced by low levels of DNA polymerase a model for chromosome fragile sites. Cell 120:587–98.

34. Iijima K, Ogata T. 2010. Construction and evaluation of self-cloning bottom-fermenting yeast with high SSU1 expression. 109:1906–1913.

35. Avram D, Leid M, Bakalinsky AT. 1999. Fzf1p of Saccharomyces cerevisiae is a positive regulator of SSU1 transcription and its first zinc finger region is required for DNA binding. 15:473–480.

36. García-Ríos E, Nuévalos M, Barrio E, Puig S, Guillamón JM. 2019. A new chromosomal rearrangement improves the adaptation of wine yeasts to sulfite. Environ Microbiol 21:1771–1781.

37. Pérez-Ortín JE, Querol A, Puig S, Barrio E. 2002. Molecular characterization of a chromosomal rearrangement involved in the adaptive evolution of yeast strains. Genome Res 12:1533–9.

38. Zimmer A, Durand C, Loira N, Durrens P, Sherman DJ, Marullo P. 2014. QTL Dissection of Lag Phase in Wine Fermentation Reveals a New Translocation Responsible for Saccharomyces cerevisiae Adaptation to Sulfite. PLOS ONE 9:e86298.

39. Mieczkowski PA, Lemoine FJ, Petes TD. 2006. Recombination between retrotransposons as a source of chromosome rearrangements in the yeast Saccharomyces cerevisiae. DNA Repair 5:1010–1020.

40. Lawrence SJ, Nicholls S, Box WG, Sbuelz R, Bealin-Kelly F, Axcell B, Smart KA. 2013. The Relationship between Yeast Cell Age, Fermenter Cone Environment, and Petite Mutant Formation in Lager Fermentations. Journal of the American Society of Brewing Chemists 71:90–96.

41. Jenkins CL, Lawrence SJ, Kennedy AI, Thurston P, Hodgson JA, Smart KA. 2018. Incidence and Formation of Petite Mutants in Lager Brewing YeastSaccharomyces Cerevisiae(Syn.S. Pastorianus) Populations. Journal of the American Society of Brewing Chemists 67:72–80.

42. Good L, Dowhanick TM, Ernandes JE, Russell I, Stewart GG. 1993. Rho− Mitochondrial Genomes and Their Influence on Adaptation to Nutrient Stress in Lager Yeast Strains. Journal of the American Society of Brewing Chemists 51:35–39.

43. Puddu F, Herzog M, Selivanova A, Wang S, Zhu J, Klein-Lavi S, Gordon M, Meirman R, Millan-Zambrano G, Ayestaran I, Salguero I, Sharan R, Li R, Kupiec M, Jackson SP. 2019. Genome architecture and stability in the Saccharomyces cerevisiae knockout collection. Nature 573:416–420.

44. De Chiara M, Barre BP, Persson K, Irizar A, Vischioni C, Khaiwal S, Stenberg S, Amadi OC, Zun G, Dobersek K, Taccioli C, Schacherer J, Petrovic U, Warringer J, Liti G. 2022. Domestication reprogrammed the budding yeast life cycle. Nat Ecol Evol 6:448–460.

45. Muenzner J, Trébulle P, Agostini F, Zauber H, Messner CB, Steger M, Kilian C, Lau K, Barthel N, Lehmann A, Textoris-Taube K, Caudal E, Egger A-S, Amari F, De Chiara M, Demichev V, Gossmann TI, Mülleder M, Liti G, Schacherer J, Selbach M, Berman J, Ralser M. 2024. Natural proteome diversity links aneuploidy tolerance to protein turnover. Nature 630:149–157.

46. Jambhekar A, Amon A. 2008. Control of Meiosis by Respiration. Current Biology 18:969–975.

47. White TJ, Bruns T, Lee S, Taylor J. 1990. 38 – Amplification and Direct Sequencing of Fungal Ribosomal RNA Genes for Phylogenetics, p 315–322. In Innis MA, Gelfand DH, Sninsky JJ, White TJ (ed), PCR Protocols 10.1016/B978-0-12-372180-8.50042-1. Academic Press, San Diego.

48. Bolger AM, Lohse M, Usadel B. 2014. Trimmomatic: a flexible trimmer for Illumina sequence data. Bioinformatics 30:2114–2120.

49. Gallone B, Steensels J, Mertens S, Dzialo MC, Gordon JL, Wauters R, Thesseling FA, Bellinazzo F, Saels V, Herrera-Malaver B, Prahl T, White C, Hutzler M, Meussdoerffer F, Malcorps P, Souffriau B, Daenen L, Baele G, Maere S, Verstrepen KJ. 2019. Interspecific hybridization facilitates niche adaptation in beer yeast. Nat Ecol Evol 3:1562–1575.

50. Scannell DR, Zill OA, Rokas A, Payen C, Dunham MJ, Eisen MB, Rine J, Johnston M, Hittinger CT. 2011. The Awesome Power of Yeast Evolutionary Genetics: New Genome Sequences and Strain Resources for the Saccharomyces sensu stricto Genus. G3 Genes|Genomes|Genetics 1:11–25.

51. Li H, Durbin R. 2009. Fast and accurate short read alignment with Burrows-Wheeler transform. Bioinformatics 25:1754–60.

52. Danecek P, Bonfield JK, Liddle J, Marshall J, Ohan V, Pollard MO, Whitwham A, Keane T, McCarthy SA, Davies RM, Li H. 2021. Twelve years of SAMtools and BCFtools. Gigascience 10.

53. Ortiz EM. 2019. vcf2phylip v2.0: convert a VCF matrix into several matrix formats for phylogenetic analysis. 10.5281/. Accessed

54. Minh BQ, Schmidt HA, Chernomor O, Schrempf D, Woodhams MD, von Haeseler A, Lanfear R. 2020. IQ-TREE 2: New Models and Efficient Methods for Phylogenetic Inference in the Genomic Era. Molecular Biology and Evolution 37:1530–1534.

55. Ram Y, Dellus-Gur E, Bibi M, Karkare K, Obolski U, Feldman MW, Cooper TF, Berman J, Hadany L. 2019. Predicting microbial growth in a mixed culture from growth curve data. Proc Natl Acad Sci U S A 116:14698–14707.

56. Schneider CA, Rasband WS, Eliceiri KW. 2012. NIH Image to ImageJ: 25 years of image analysis. Nature Methods 9:671–675.

57. Zupan J, Avbelj M, Butinar B, Kosel J, Sergan M, Raspor P. 2013. Monitoring of quorum-sensing molecules during minifermentation studies in wine yeast. J Agric Food Chem 61:2496–505.

58. Yue JX, Li J, Aigrain L, Hallin J, Persson K, Oliver K, Bergstrom A, Coupland P, Warringer J, Lagomarsino MC, Fischer G, Durbin R, Liti G. 2017. Contrasting evolutionary genome dynamics between domesticated and wild yeasts. Nat Genet 49:913–924.

59. Weiß CL, Pais M, Cano LM, Kamoun S, Burbano HA. 2018. nQuire: a statistical framework for ploidy estimation using next generation sequencing. BMC Bioinformatics 19:122.

60. Smit AFA, Hubley R, Green P. RepeatMasker http://repeatmasker.org. Accessed

61. Li HJaG. 2013. Aligning sequence reads, clone sequences and assembly contigs with BWA-MEM.

62. van der Auwera G, O’Connor BD. 2020. Genomics in the Cloud: Using Docker, GATK, and WDL in Terra. O’Reilly Media, Incorporated.

63. Van der Auwera GA, Carneiro MO, Hartl C, Poplin R, del Angel G, Levy-Moonshine A, Jordan T, Shakir K, Roazen D, Thibault J, Banks E, Garimella KV, Altshuler D, Gabriel S, DePristo MA. 2013. From FastQ Data to High-Confidence Variant Calls: The Genome Analysis Toolkit Best Practices Pipeline. 43:11.10.1–11.10.33.

64. Robinson JT, Thorvaldsdóttir H, Winckler W, Guttman M, Lander ES, Getz G, Mesirov JP. 2011. Integrative genomics viewer. Nature Biotechnology 29:24–26.

65. D’Angiolo M, De Chiara M, Yue JX, Irizar A, Stenberg S, Persson K, Llored A, Barre B, Schacherer J, Marangoni R, Gilson E, Warringer J, Liti G. 2020. A yeast living ancestor reveals the origin of genomic introgressions. Nature 587:420–425.

66. Cingolani P, Patel VM, Coon M, Nguyen T, Land SJ, Ruden DM, Lu X. 2012. Using Drosophila melanogaster as a Model for Genotoxic Chemical Mutational Studies with a New Program, SnpSift. Front Genet 3:35.

67. Suvakov M, Panda A, Diesh C, Holmes I, Abyzov A. 2021. CNVpytor: a tool for copy number variation detection and analysis from read depth and allele imbalance in whole-genome sequencing. GigaScience 10.

68. O’Donnell S, Yue J-X, Saada OA, Agier N, Caradec C, Cokelaer T, De Chiara M, Delmas S, Dutreux F, Fournier T, Friedrich A, Kornobis E, Li J, Miao Z, Tattini L, Schacherer J, Liti G, Fischer G. 2023. Telomere-to-telomere assemblies of 142 strains characterize the genome structural landscape in Saccharomyces cerevisiae. Nature Genetics 55:1390–1399.

69. De Chiara M, Friedrich A, Barre B, Breitenbach M, Schacherer J, Liti G. 2020. Discordant evolution of mitochondrial and nuclear yeast genomes at population level. BMC Biol 18:49.

70. Fernandez-Ricaud L, Kourtchenko O, Zackrisson M, Warringer J, Blomberg A. 2016. PRECOG: a tool for automated extraction and visualization of fitness components in microbial growth phenomics. BMC Bioinformatics 17:249.

